# A genomics approach reveals the global genetic polymorphism, structure and functional diversity of ten accessions of the marine model diatom *Phaeodactylum tricornutum*

**DOI:** 10.1101/176008

**Authors:** Achal Rastogi, FRJ Vieira, Anne-Flore Deton-Cabanillas, Alaguraj Veluchamy, Catherine Cantrel, Gaohong Wang, Pieter Vanormelingen, Chris Bowler, Gwenael Piganeau, Hanhua Hu, Leila Tirichine

**Affiliations:** Institut de biologie de l’Ecole normale supérieure (IBENS), Ecole normale supérieure, CNRS, INSERM, PSL Université Paris 75005 Paris, France; Key Laboratory of Algal Biology, Institute of Hydrobiology, Chinese Academy of Sciences, Wuhan, 430072 China; Ghent University, Department of Biology, Research Group Protistology and Aquatic Ecology Krijgslaan 281/S8 9000 Gent, Belgium; Sorbonne Universités, UPMC Univ Paris 06, CNRS, Biologie Intégrative des Organismes Marins (BIOM), Observatoire Océanologique, F-66650 Banyuls/Mer, France

## Abstract

Diatoms emerged in the Mesozoic period and presently constitute one of the main primary producers in the world’s ocean and are of a major economic importance. In the current study, using whole genome sequencing of ten accessions of the model diatom *Phaeodactylum tricornutum*, sampled at broad geospatial and temporal scales, we draw a comprehensive landscape of the genomic diversity within the species. We describe strong genetic subdivisions of the accessions into four genetic clades (A-D) with constituent populations of each clade possessing a conserved genetic and functional makeup, likely a consequence of the limited dispersal of *P. tricornutum* in the open ocean. We further suggest dominance of asexual reproduction across all the populations, as implied by high linkage disequilibrium. Finally, we show limited yet compelling signatures of genetic and functional convergence inducing changes in the selection pressure on many genes and metabolic pathways. We propose these findings to have significant implications for understanding the genetic structure of diatom populations in nature and provide a framework to assess the genomic underpinnings of their ecological success and impact on aquatic ecosystems where they play a major role. Our work provides valuable resources for functional genomics and for exploiting the biotechnological potential of this model diatom species.

## Introduction

Diatoms are unicellular predominantly diploid and photosynthetic eukaryotes. They belong to a big group of heterokonts, constituent of chromalveolate (SAR group) which are believed to be derived from serial endosymbiosis combining genes from green and red algae predecessors and further diversified via horizontal gene transfer from a wide range of prokaryotes [1–3]. Ehrenberg [4] first discovered diatoms in the 19th century in dust samples collected by Charles Darwin in the Azores. According to the earliest fossil records, they are believed to be in existence since at least 190 million years [5]. In nature, most diatoms likely live in obligate relationships with bacteria [6] but many, like *Phaeodactylum tricornutum*, can be propagated in axenic conditions. In spite of its low abundance in the open ocean [7], *P. tricornutum* is extensively used as a model to study and characterize diatom metabolism, and to understand diatom evolution [1, 8–12].

*P. tricornutum* is a coastal diatom found under highly unstable environments like estuaries and rock-pools. Although it has never been reported to undergo sexual reproduction, factors such as sensitivity to many nonspecific abiotic components and the general lack of knowledge on sexual reproduction in this species [13–15] limit our ability to constrain the sexual cycle of these organisms. Since the discovery of *P. tricornutum* by Bohlin in 1897 and the characterization of different morphologies, denoted fusiform, triradiate, oval, round and cruciform, 10 isolates from 9 different geographic locations (sea shores, estuaries, rock pools, tidal creeks) around the world, from sub-polar to tropical latitudes, have been accessioned (Fig. S1) [well described in [16]. These accessions have been collected within the time frame of approximately one century, from 1908 (Plymouth isolate, Pt2/3) to 2000 (Dalian isolate, Pt10) (Fig. S1) [16]. All the isolates have been maintained either axenically or with native bacterial populations in different stock centers and have been cryopreserved after isolation. Previous studies have reported distinct functional behaviors of different accessions as adaptive responses to various environmental cues [17–20], but very little is known about their genetic diversity. However, based on sequence similarity of the ITS2 region within the 28S rDNA repeat sequence, the accessions can be divided into four genotypes (Genotype A: Pt1, Pt2, Pt3 and Pt9; Genotype B: Pt4; Genotype C: Pt5 and Pt10; Genotype D: Pt6, Pt7 and Pt8), with genotypes B and C being the most distant [16]. *P. tricornutum* is among the few diatom species with a whole genome sequence available to the community [21], and the only diatom for which extensive state-of-the-art functional and molecular tools have been developed over the past few decades [22–35]. These resources have advanced *P. tricornutum* as a model diatom species and provided a firm platform for future genome-wide structural and functional studies.

The accumulated effects of diverse evolutionary forces such as recombination, mutation, and selection have been found to dictate the structure and diversity of genomes in a wide range of species [36–39]. The existence of genomic diversity within a species reflects its potential to adapt to a changing environment. Exploring the genomic diversity within a species not only provides information about its evolution, it also offers opportunities to understand the role of various biotic and abiotic interactions in structuring a genome [40]. Such studies in diatoms are rare and estimates of genetic diversity within diatom populations are mostly inferred using microsatellite-based genotyping approaches [41–43]. Although these techniques have revealed a wealth of information about diatom evolution, their dispersal and reproductive physiology [40], additional insights can be obtained using state-of-the-art whole genome comparative analysis techniques [43]. Deciphering the standing genomic variation of *P. tricornutum* across different accession populations, sampled at broad geospatial scale, is an important first step to assess the role of various evolutionary forces in regulating the adaptive capacities of diatoms in general (e.g.[44]). To understand the underlying genomic diversity within different accessions of *P. tricornutum* and to establish the functional implications of such diversity, we performed deep whole genome sequencing of the 10 most studied accessions, referred to as Pt1 to Pt10 [16, 19, 45]. We present a genome-wide diversity map of geographically distant *P. tricornutum* accessions, describing a stable genetic structure in the environment. This work further provides the community with whole genome sequences of the accessions, which will be a valuable genetic resource for functional studies of accession-specific ecological traits in the future.

## Results

### Reference-assisted assembly reveals low nucleotide diversity across multiple accessions of *P. tricornutum*

We sequenced the whole genomes of ten accessions of *P. tricornutum* using Illumina HiSeq 2000, and performed a reference-based assembly using the genome sequence of the reference strain Pt1 8.6 [1]. Across all accessions, the percentage of sequence reads mapped on the reference genome ranged between ~65% to ~80% (Table 1), with an alignment depth ranging between 26X and 162X, covering 92% to 98% of the reference genome (Table 1). Many regions on the reference genome that are observed as being unmapped by reads from individual ecotypes are annotated as rich in transposable elements (TEs) (Fig. S2). At >90% identity, the repeated proportion of unmapped reads varies between ~38% (Pt1) and 75% (Pt4).

**Table 1.** Reference-assisted mapping statistics. The table summarizes the origin and year of sampling of each accession of *P. tricornutum* along with the number of total reads mapped on the reference. Average depth (X=average number of reads aligned on each base covered across the entire genome) was estimated using the number of mapped read pairs and the horizontal coverage (aka. coverage breadth) across the whole genome.

| Library Name | Accession number/axenicity | Origin | Year of Isolation | Mapped Read-Pairs | % Mapped Read-Pairs | Alignment Depth (X) | Genome Coverage (%) |
| --- | --- | --- | --- | --- | --- | --- | --- |
| Pt1 | CCMP2561 (axenic) | Blackpool, UK | 1956 | 3, 642,044 | 79.41 | 26.5 | 98.0 |
| Pt2 | CCMP2557 (xenic) | Plymouth, UK | Prior to 1910 | 6, 016,241 | 78.23 | 43.8 | 98.0 |
| Pt3 | CCMP2558 (axenic) | Plymouth, UK | 1930s | 6, 373,591 | 65.62 | 46.4 | 98.3 |
| Pt4 | CCMP2559 (xenic) | Finland | 1951 | 15, 583,665 | 67.31 | 113.5 | 94.0 |
| Pt5 | CCMP630 (axenic) | West Dennis, MA, USA | 1972 | 5, 346,009 | 75.50 | 38.9 | 93.2 |
| Pt6 | CCMP631 (xenic) | MA, USA | 1956 | 3, 922,830 | 64.50 | 28.5 | 94.1 |
| Pt7 | CCMP1327 (axenic) | Long Island, NY, USA | 1952 | 4, 937,516 | 67.30 | 35.9 | 94.9 |
| Pt8 | CCMP2560 (xenic) | Vancouver, Canada | 1987 | 22, 235,170 | 78.36 | 162.1 | 94.4 |
| Pt9 | CCMP633 (axenic) | Guam, Micronesia | 1981 | 7, 551,099 | 74.68 | 55.2 | 97.5 |
| Pt10 | CCMP2928 (axenic) | Dalian, China | 2000 | 5, 436,057 | 72.59 | 39.6 | 92.1 |

Following the assembly, we performed variant calling using Genome Analysis Toolkit [46] and discovered 462,514 (depth >= 4x) single nucleotide polymorphisms (SNPs) including ~25% singleton sites, 573 insertions (of varying lengths from 1 bp to 312 bp) and 1,801 deletions (of lengths from 1 bp to 400 bp) (Fig. 1A), across all the accessions. The spectrum of SNPs across all the accessions further reveals a higher rate of transitions (Ts) over transversions (Tv) (Ts/Tv = 1.6). In total, compared to the reference alleles from Pt1.8.6, six possible types of single nucleotide changes could be distinguished, among which G:C -> A:T and A:T -> G:C accounted for more than ∼60% of the observed mutations (Fig. S3A). Further, most SNPs and INDELs (insertions and deletions) are shared between different accessions, except for Pt4, which possesses the highest proportion of specific SNPs (~35%) and INDELs (~75%) (Fig. 1B). Interestingly, we found that most of the SNPs are heterozygous, and the proportion of heterozygous variants across all the accessions varies between ~45% (in Pt5 and Pt10) to ~98% (in Pt1, Pt2 and Pt3) (Fig. 1C). Most of the variant alleles in the accessions with high proportions of heterozygous variants were further found to be significantly deviated from Hardy-Weinberg equilibrium (HWE) (chi-square test, P-value < 0.05) (Fig. 1C), possibly linked to prolonged asexual reproduction [47]. Surprisingly, despite significant differences in the proportion of heterozygote variant alleles between the accessions, which ranges between 45% to 98%, the average pairwise synonymous nucleotide diversity (π_S_) estimated from genes with callable sites across all the accessions is 0.007 per synonymous site. This indicates that any two homologous sequences taken at random across different populations will on average differ by only ∼0.7% on synonymous positions. The non-synonymous pairwise diversity (π_N_) over the same genes is 0.003, consistent with an excess of non-synonymous mutations being deleterious. An average non-synonymous (N) to synonymous (S) variant ratio (π_N_ / π_S_) was estimated to be ~0.43, which is higher than in the *Ostreococcus tauri*, π_N_ / π_S_ = 0.2 [48]. Since π_N_ / π_S_ is negatively correlated with the effective population size, which is the number of distinct clones in unicellular organisms[49], this could reflect either a lower census population size in *P. tricornutum*, and/or a lower rate of sexual reproduction. Linkage disequilibrium (LD) analysis using only homozygous SNP sites revealed, on average, high LD (>0.7) over pairs of variations, genome wide (Fig. S3B). Further, based on the difference in the allelic frequencies of the SNPs, the pairwise *Fst* between the populations ranges from ∼0.005 (between Pt1 and Pt3) to ∼0.4 (between Pt4 and Pt10) (Fig. 1D). Considering *Fst* as a measure of genetic differentiation or structuring between the populations, the ten *P. tricornutum* accessions can be clustered into 4 genetic groups/clades with Pt1, Pt2, Pt3 and Pt9 in clade A; Pt4 in clade B; Pt5, Pt10 in clade C; and Pt6, Pt7, Pt8 in clade D, reflecting low intra-group Fst (∼0.02) and high inter-group Fst (0.2 – 0.4) (Fig. 1D).

**Figure 1.**
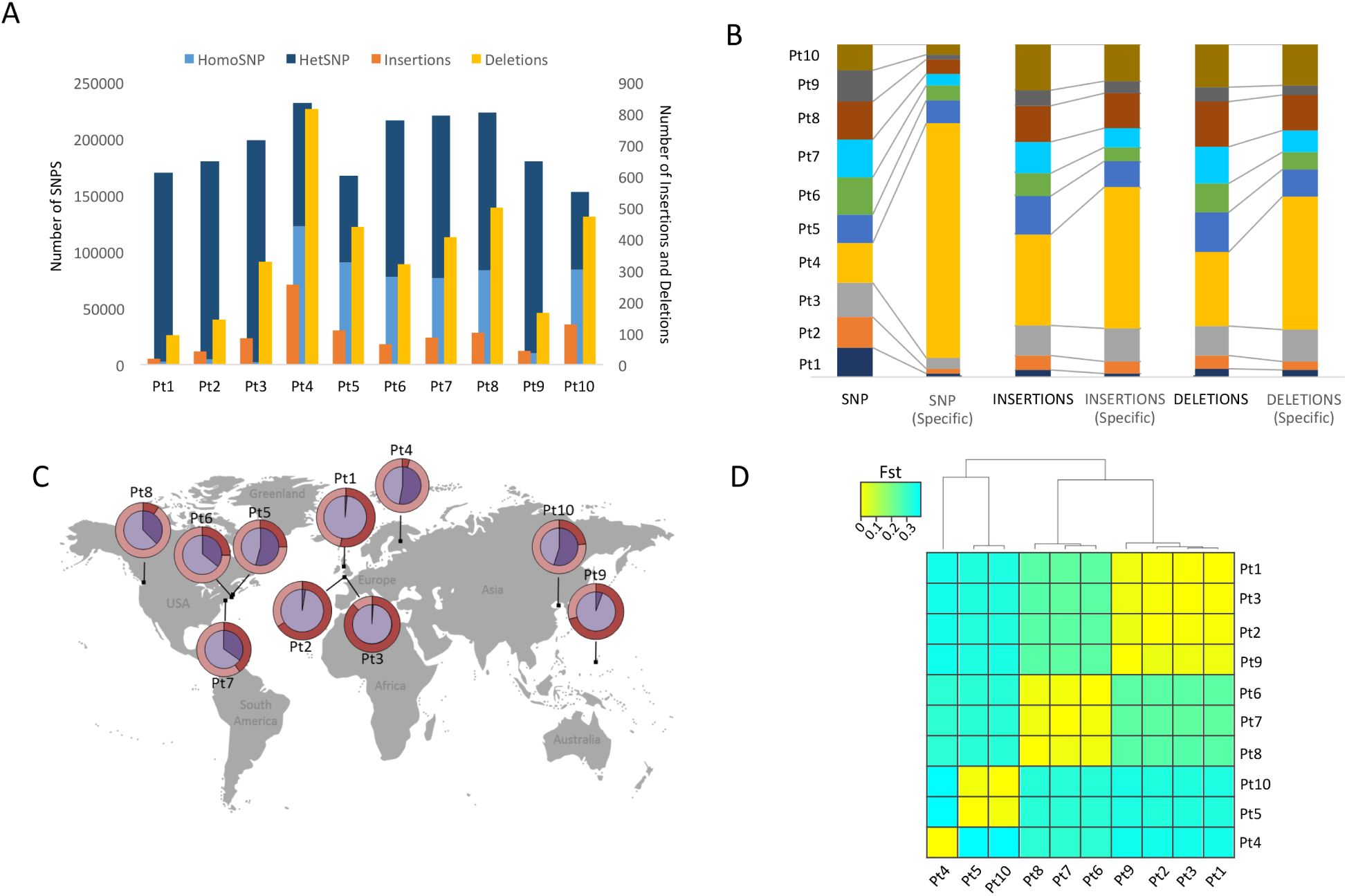
Genetic diversity between *P. tricornutum* accessions. (A) The bar plot represents total number of discovered SNPs, with the proportion of heterozygous SNPs (dark blue) and homozygous SNPs (light blue), INSERTIONS (orange) and DELETIONS (yellow) in each accession compared to the reference genome. (B) The stack bar plot represents the proportion of total vs specific polymorphic variant sites, including SNPs, insertions and deletions (from left to right, respectively) across all the accessions. (C) The world map indicates proportion of heterozygous (dark violet) and homozygous SNPs (violet) in each accession, represented as pie charts. The outer ring represents the proportion of variant alleles being significantly deviated from HWE (deep red). (D) The heat-map shows the genetic differentiation or association between all possible pairs of accessions. The colors indicate FST values, which range from 0.02 to 0.4, with a color gradient from yellow to green, respectively. Values closer to 0 signify close genetic makeup and values closer to 1 indicate strong genetic structuring between the populations.

### Four genetic clades of *P. tricornutum*

With the exception of Pt4, where we found the maximum number of variant alleles to be accession-specific, most of the variant alleles are shared between at least two accessions, indicating close genetic relatedness (Fig. 1B). Therefore, in order to cluster the accessions based on the genome structure shared among them, we used Bayesian clustering approach by applying *Markov Chain Monte Carlo* (MCMC) estimations, programmed within the ADMIXTURE software [50]. Based on the allelic composition of the ten genomes, six genomic clusters (K=6) can be formed, which is distributed within each accession genome in different proportions (represented by pie-chart colors) (Fig. 2A). Further, depending on the pattern (both qualitative and quantitative) of distributed genomic clusters across different accession genomes, the ten accessions revealed four genetic clusters with Pt1, Pt2, Pt3, and Pt9 in one, Pt4 in a second, Pt5, Pt10 in a third, and Pt6, Pt7, Pt8 in a fourth cluster (Fig. 2B). These clusters (Fig. 2B) are in broad agreement with *Fst*-based genetic clades (Fig. 1D), phylogenetic clusters inferred using ribosomal marker genes (18S (Fig. S4A), and ITS2 (Fig S4B)), as also reported previously [16], and at whole genome scale (this study) as inferred by a phylogenetic tree generated using maximum likelihood algorithm based on all (Fig. 2C) and only homozygous polymorphic sites (SNVs and INDELS) (Fig. S4C).

**Figure 2:**
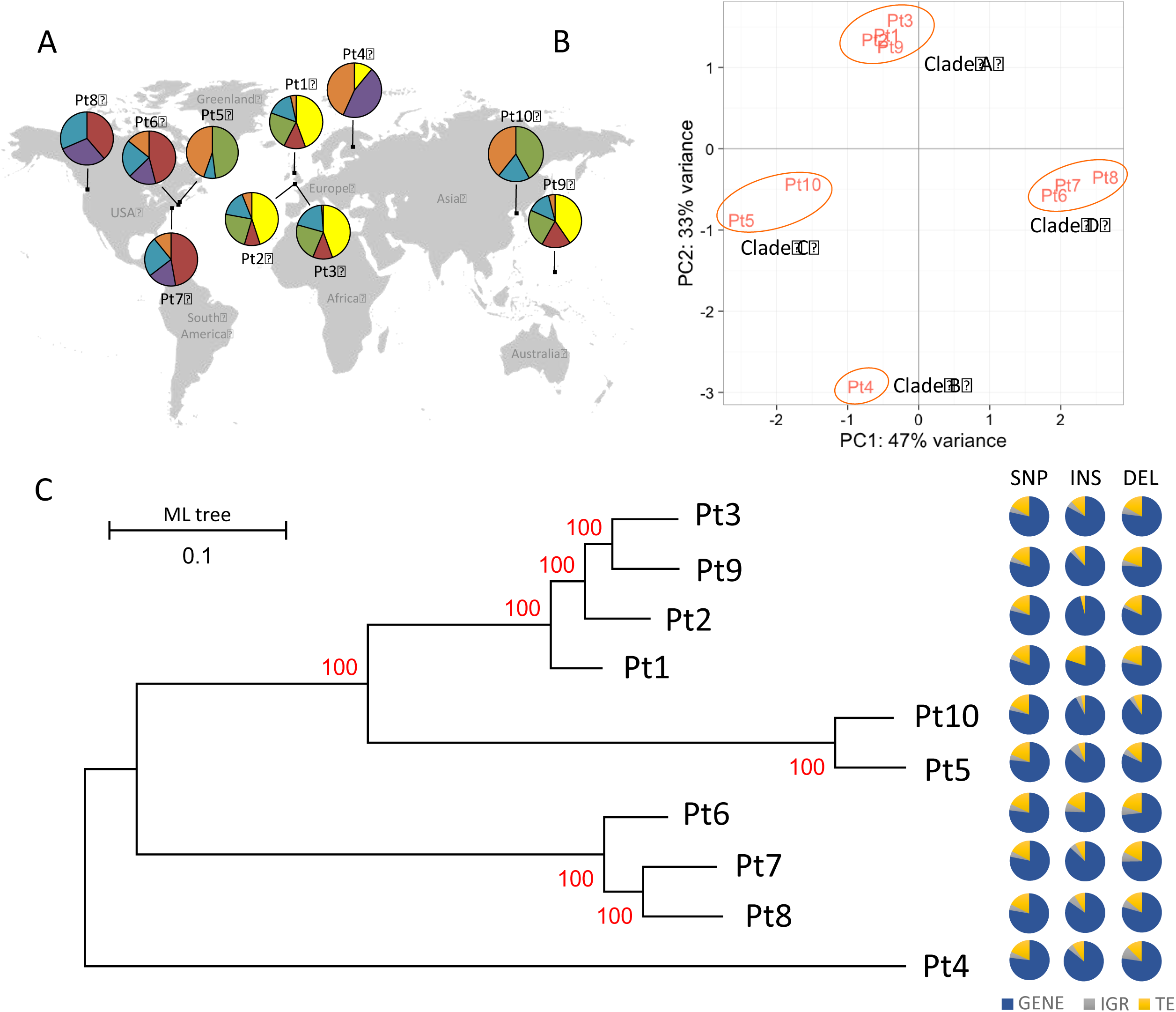
Clustering of *P. tricornutum* accessions. (A) Principal component analysis (PCA) showing the distribution of the ten accessions based on their shared genome structure, revealing four genetic clusters referred to as clades A, B, C, and D. (B) Pie charts showing the genetic makeup of the genomes of each accession. Based on the allelic composition of the ten genomes, six genomic clusters (K=6) are formed, which are distributed across individual accession genome in different proportions (represented by 6 different colors in the pie-chart). (C) Phylogenetic association of the accessions based on 468,188 genome-wide polymorphic sites (including SNP and INDELS) using a maximum likelihood approach. The numbers on the branches indicate the bootstrap values. Pie charts adjacent to each node of the whole genome tree correspond to the proportion of SNPs and INDELs over all functional features of the genome; GENEs (blue), TEs (yellow), IGRs (Intergenic Regions, represented in grey).

Further sequential assessment of the 18S and ITS2 rDNA gene sequences across different clades indicated the presence of multiple variations, including both heterozygous and homozygous variant alleles (Fig. S4D and S4E). Because the ribosomal DNA region including 18S and ITS2 is highly repetitive, which is on average ~4 times more than non-ribosomal genes (Fig. 3A), these differences can be understood as intra-genomic variations within the genome. However, taxonomists and ecologists use differences within 18S gene sequences as a measure of species assignation and to estimate species delineation [7]. This latter practice has been previously shown to be very conservative as no differences in the 18S gene were found between reproductively isolated species [51]. Alternatively, the possibility of sub-populations or cryptic populations cannot be ignored, as previously reported in planktonic foraminifers [52] and coccolithophores [53].

**Figure 3.**
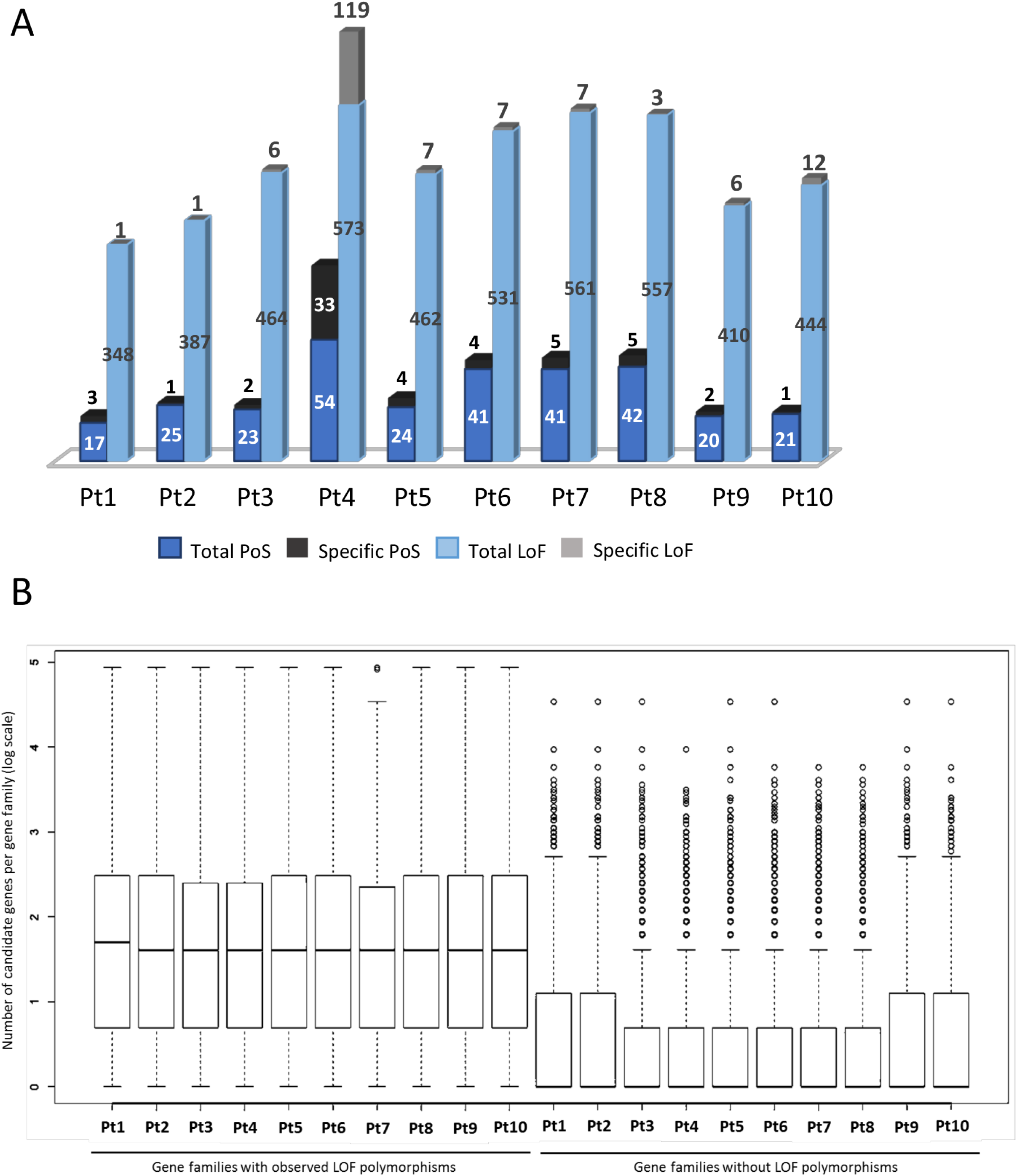
Large structural variations within accessions. (A) The heat-map displays the fold-change (FC) of read depth between each reference gene and median of read depth of all the reference genes, within each accession. Using Z-score as a measure of normalized read depth, log2 fold change (FC) is calculated as a ratio of Z-score per gene to the average normalized read depth of all the genes per accession. A blue to red color gradient in the heat-map represents low to high log2FC. From all the accessions only those genes are plotted where log2FC is more than 2 in at least one of the accessions and are considered to exhibit copy number variation (CNV). (B) The bar plots represent the total and specific numbers of genes, denoted on Y-axis, that exhibit a loss or multiple copies (CNV) within one or more accessions. (C) The bar plots represent the number of total-and accession-specific TEs exhibiting CNVs across one or more accessions. (D) With similar principle aesthetics as panel (A) of this figure, the heat-map shows the patterns of log2FC only across all the accessions of those TEs exhibiting CNV in at least one of the ten accessions studied.

We examined the possible presence of sub-populations on 18S gene heterozygosity in some of the accessions. We confirmed the expression of all the heterozygous alleles within the 18S rDNA gene using whole genome and total-RNA sequencing of a monoclonal culture (propagated from a single cell) from Pt8 (constituent of Clade D) and Pt3 (constituent of Clade A) population, referred to as Pt8Tc and Pt3Ov (Fig. S4D), respectively. This experiment indicates that the cultures (Pt1-Pt10) are a single population with no or undetectable heterogeneity.

Next, concerning the observed polymorphisms within the 18S ribosomal marker gene, we investigated whether the four clades can be considered as different species. We looked for the existence of compensatory base changes (CBCs) within secondary structures of the ITS2 gene between all pairs of accessions. The presence of CBCs within ITS2 has been recently suggested to account for reproductive isolation in multiple plant species [54] and between diatom species [55, 56]. By comparing the ITS2 secondary structure from all the accessions, we did not find any CBCs between any given pair of accessions (Fig. S5). As a control, we compared the ITS2 secondary structure of all the *P. tricornutum* accessions with the ITS2 sequences of other diatom species (*Cyclotella meneghiniana*, *Pseudo-nitzschia delicatissima*, *Pseudo-nitzschia multiseries*, *Fragilariopsis cylindrus*) that have significant degrees of evolutionary divergence as depicted previously using multiple molecular marker genes [21, 57], and found multiple CBCs in them (Fig. S5).

### Close genetic relatedness depicted by large structural genomic variations among accessions

Next, using a normalized measure of read depth (see Materials and Methods), we found that 259 and 590 genes, representing ~2% and ~5% of the total gene content, respectively, have been lost or exhibit copy number variation (CNV), across the ten accessions (Fig. 3A, 3B) (File S1). Multiple randomly chosen loci were also validated by PCR for their loss from certain accessions compared to the reference strain Pt1 8.6 (Fig. S6). Compared to the reference, approximately 70% of the genes that are either lost or show CNV are shared among multiple accessions with an exception of Pt10, which displays the maximum number of lost genes and accession-specific genes exhibiting CNV (Fig. 3B). In addition, we detected 207 TEs (~6% of the total annotated TEs) (File S2) showing CNVs across one or more accessions (Fig. 3C, 3D), 80% of which are shared among two or more accessions, with Pt10 again possessing the maximum number of accession-specific TEs exhibiting CNVs (Fig. 3C). Not surprisingly, across all the accessions, class I-type TEs, which undergo transposition via a copy-and-paste mechanism, show more variation in the estimated number of copies than class II-type TEs (Fig. 3D, S7) that are transposed by a cut-and-paste mechanism. Euclidean distance estimated between accessions, based on the variation in the number of copies of different genes and TEs displaying CNVs, followed by hierarchical clustering, depicted three genetic clusters: Pt1, Pt2, Pt3, Pt9 in cluster1; Pt5, Pt10 in cluster 2, and Pt4, Pt6, Pt7, Pt8 in cluster 3 (Fig. 3A, 3D). These clusters are in broad agreement with the ones described by *Fst* and indicate the closer genetic makeup between accessions within a cluster than between the clusters. Further, biological processes can only be traced for ~40% of the genes exhibiting accession-specific CNVs. Among all the enriched biological processes (chi-square test, P<0.01) (File S1), a gene associated to nitrate assimilation (Phatr3_EG02286) is observed to have higher copy number specifically in Pt4. Likewise, each accession can be characterized by specific genetic features, represented by ~0.3% to ~28% accession-specific CNVs (Fig. 3B), possibly linked to the explicit functional behavior of some accessions in response to various environmental cues, as reported previously [17–19].

### Evolutionary adaptation in *P. tricornutum* clades

Species are under continuous pressure to adapt to a changing environment over time. We therefore wanted to understand the functional consequences of the genetic diversity between the accessions. Localization of the polymorphic sites over genomic features (genes, TEs, and intergenic regions) revealed highest rate of variation within genes (Fig. 2C), specifically on exons, and was consistent across all the studied accessions. We further identified genes within different phylogenetic clades experiencing different selection pressure based on lowest and highest π_N_ / π_S_ ratios. Across all the accessions, 241 genes displaying π_N_ / π_S_ >1 and a higher frequency of non-synonymous to synonymous polymorphism, as expected under balancing selection (BS) [58] (File S3). Furthermore, across all the accessions, 128 genes exhibit a signature of relaxed selection (RS) with accumulation of NS polymorphisms, among which 47% are specific to one or other clades (Fig. 4A). In addition, many genes (902) were found to have loss-of-function (LoF hereafter) variant alleles (Fig. 4A), including frame-shift mutations and mutations leading to theoretical start/stop codon loss and/or gain.

**Figure 4.**
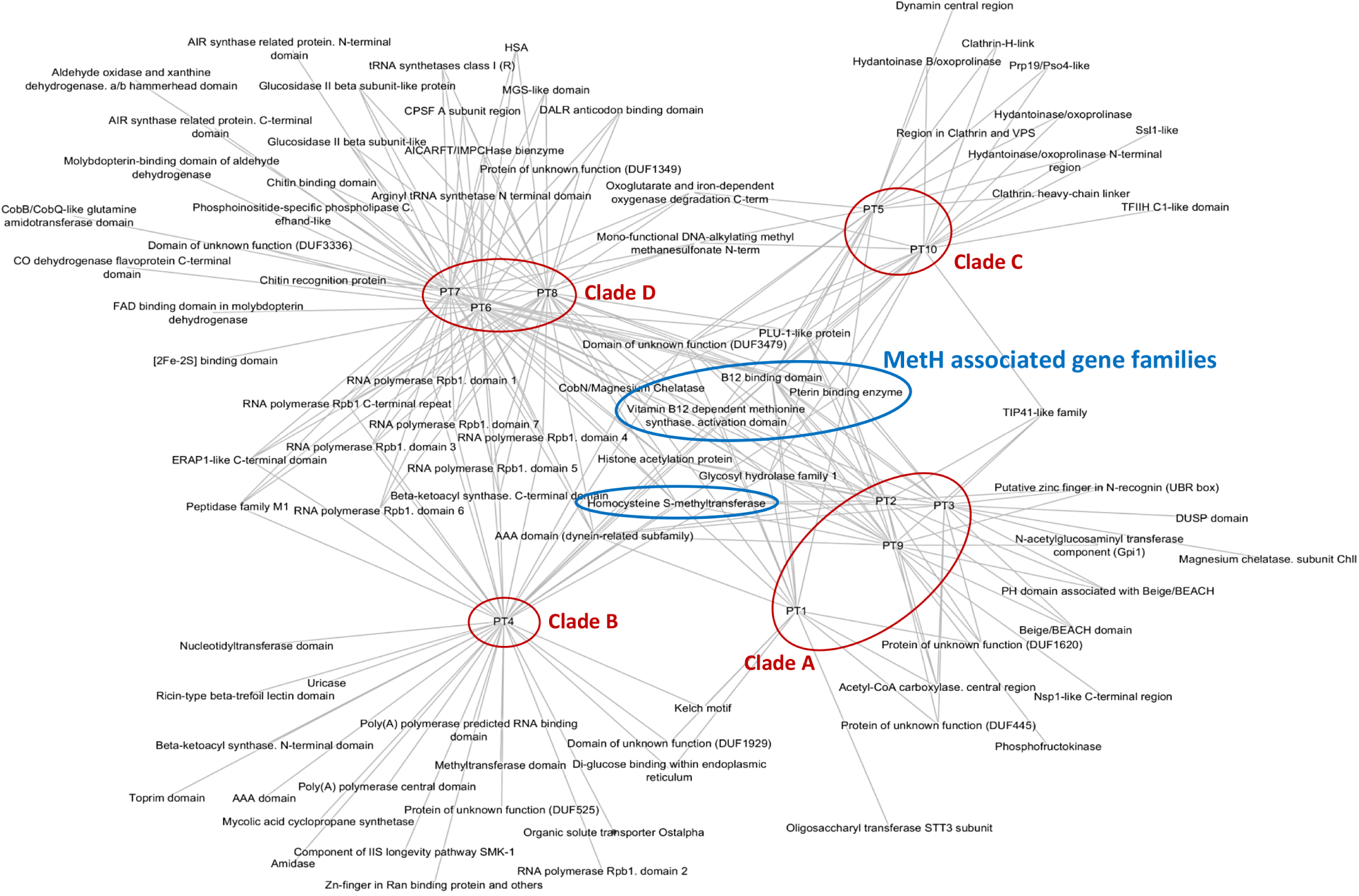
Evolutionary and functional consequences of polymorphisms. (A) The bar plot represents total and specific numbers of genes that are subject to positive selection, or experiencing loss-of-function (LoF) mutations. For each category, the accessions are plotted as stack plots with total and specific numbers of genes. Numbers of genes in each category are indicated. (B) The box plot represents the number of gene families affected by loss-of-function (LoF) mutations and suggests a bias of such mutations on the genes belonging to large gene families. Y-axis represents, as log scale, the number of genes in the gene families vs those that are not affected by LoF mutations.

Based on the presence of functional domains (Pfam domains), all *P. tricornutum* annotated genes [63] were grouped into 3,020 gene families. These families can be as large as the reverse transcriptase gene family, which is highly abundant in marine plankton [64], representing 149 candidate genes having reverse transcriptase domains, or as small as families that constitute single gene candidates. Across all the accessions, we observed that most genes experiencing LoF mutations belong to large gene families (Fig. 4B). This is consistent with a previous observation of the existence of functional redundancy in gene families as a balancing mechanism for null mutations in yeast [65]. Therefore, to estimate an unbiased effect of any evolutionary pressure (eg. LoF allele mutations) on different gene families, we calculated a ratio, termed the effect ratio (EfR, see Materials and Methods), which normalizes that if any gene family has enough candidates to buffer the effect on some genes influencing evolutionary pressure, it will be considered as being less affected compared to those for which all or most of the constituents are under selection pressure. From this analysis, each genetic clade displayed a specific set of gene families as being under selection (Fig. 5). Functional enrichment of constrained genes revealed enrichment of (1) AAA family proteins that often perform chaperone like functions that assist in the assembly or disassembly of proteins complexes, protein transport and degradation as well as other functions such as replication, recombination, repair and transcription [66], (2) tetratricopeptide-like repeats known for their role in a variety of biological processes, such as cell cycle regulation, organelle targeting and protein import, vesicle fusion and biomineralization [67]. A redox class of enzymes are common to both groups of genes and a significant proportion of unknown function proteins is found in the group of genes under balancing selection (File S3).

**Figure 5.**
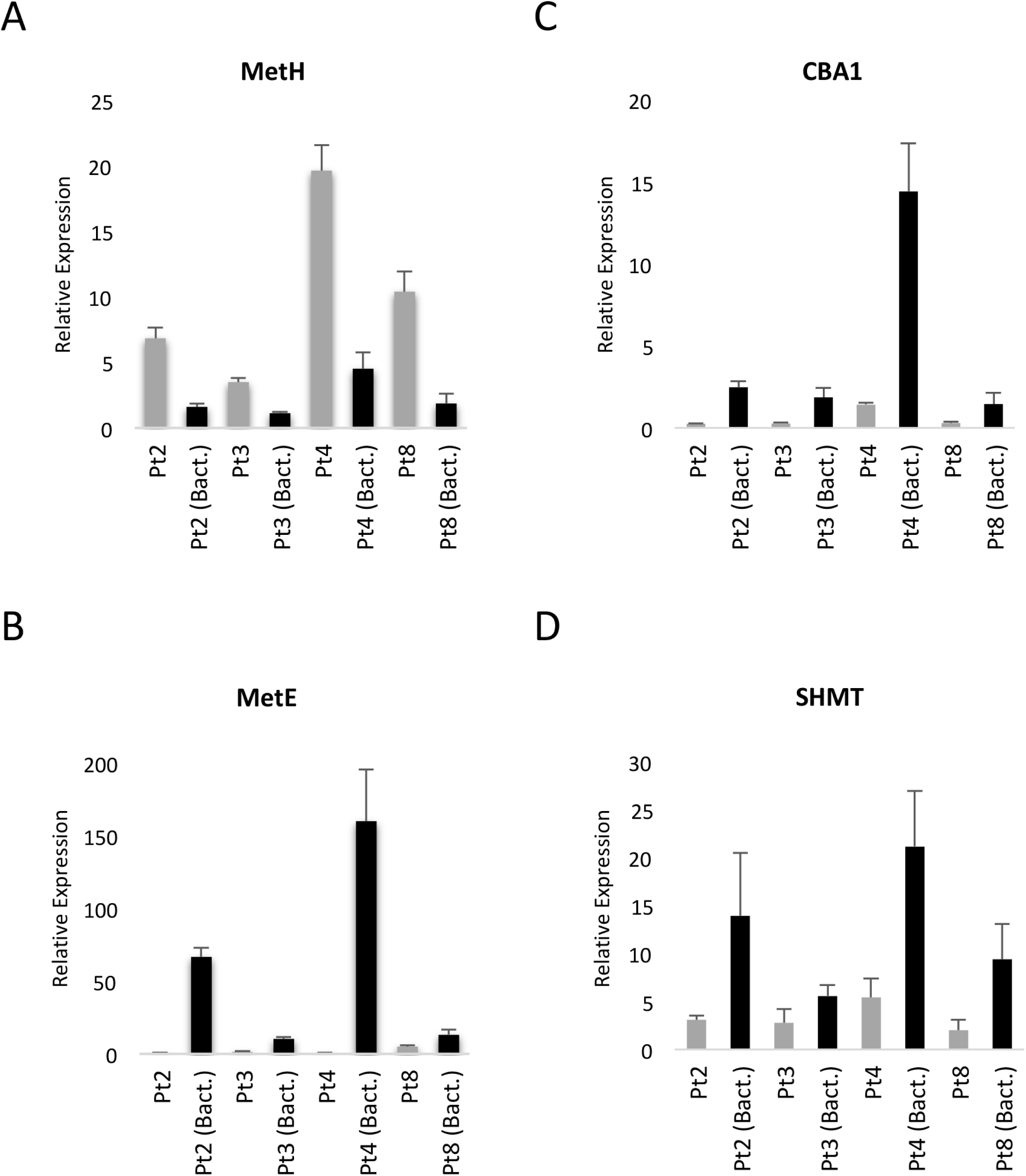
Relaxed selection within each genetic clade. Based on the EfR metric, the network displays highly affected gene families experiencing balancing selection. Gene families associated with *MetH* gene in all the accessions are indicated within the blue circles. The red circles group individual accessions as clades.

Finally, considering all pairwise correlated gene families exhibiting similar selection signals, measured using EfR among the 10 accessions, we used hierarchical clustering to examine the functional closeness of accessions with one another (See methods). Consistent with the population structure, accessions within individual clades are more closely related than the accessions belonging to other clades (Fig. S8A and S8B), suggesting variation in functional relatedness between different proposed phylogenetic clades.

### Selection of *MetH* facilitated methionine biosynthesis over *MetE*

Apart from the clade-specific genes that are under high selection pressure as depicted by high rate of N/S divergence when compared to the reference, a group of gene families associated with methionine biosynthesis *(MetH*, Phatr3_J23399) was also observed to accumulate non synonymous polymorphisms in all the accessions. In *P. tricornutum*, *MetE* (cobalamin-independent methionine synthase) and *MetH* (cobalamin-dependent methionine synthase) are known to catalyze conversion of homocysteine to methionine in the presence of symbiotic bacteria and vitamin B12 in the growth media, respectively. Previous reports have suggested that growing axenic cultures in conditions of high cobalamin (vitamin B12) availability results in repression of MetE, leading to its loss of function and high expression of the MetH gene in *P. tricornutum* and *C. reinhardtii* [68–70]. In accordance with these results, we observed a high expression of *MetH* in axenically grown laboratory cultures (Fig. 6A) compared to its expression in cells cultured with their natural co-habitant bacteria. However, we were not able to trace any significant signature for the loss of *MetE* gene although its expression is significantly lower in axenic cobalamin-containing cultures (Fig. 6B). Similar observations were obtained for *CBA1* and *SHMT* genes (Fig. 6C and 6D), which under cobalamin scarcity enhance cobalamin acquisition and manage reduced methionine synthase activity, respectively [69].

**Figure 6.**
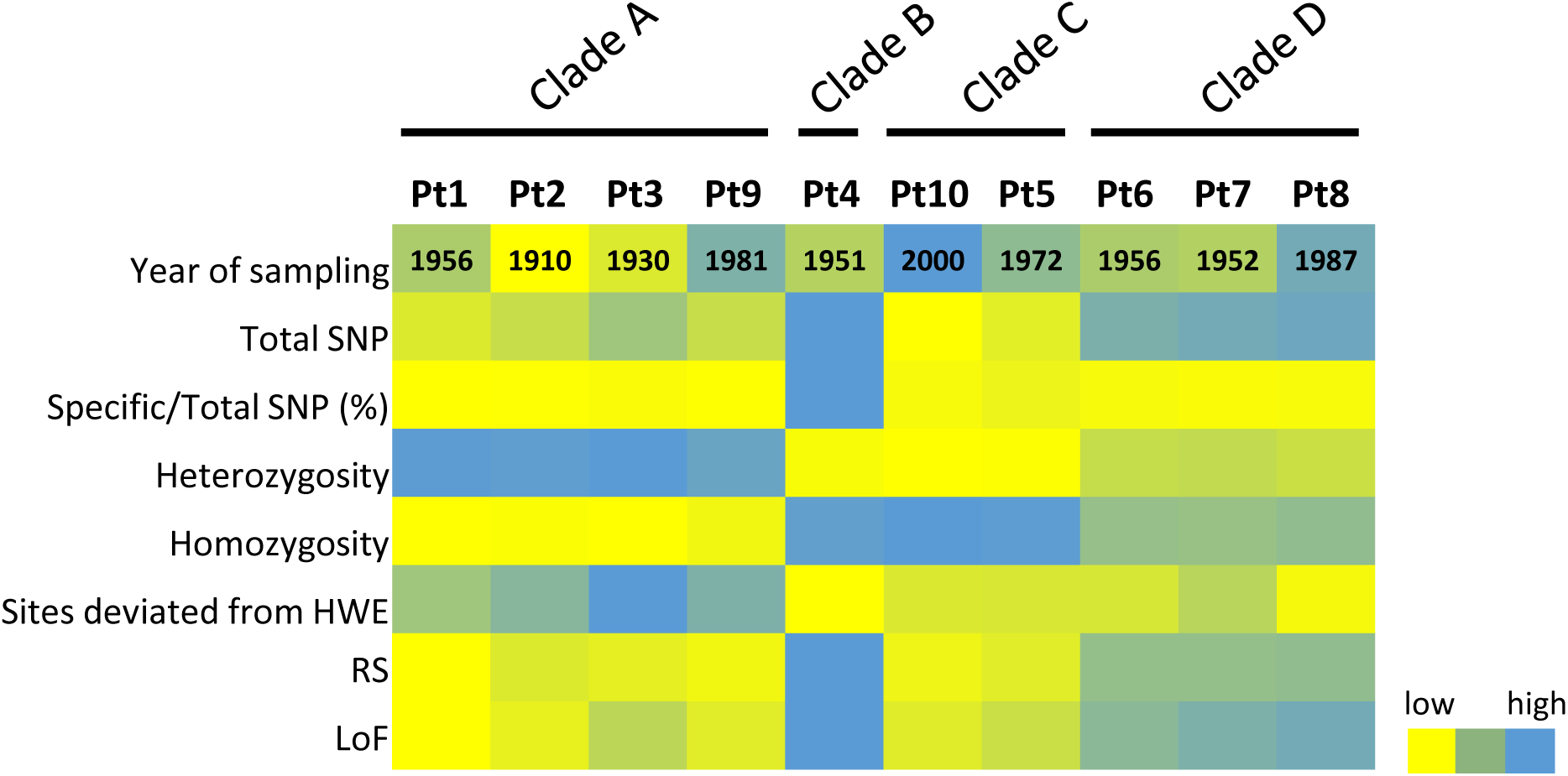
Selection of MetH-facilitated methionine biosynthesis over MetE. The bar plots represent relative expression of (A) *MetH*, (B) *MetE*, (C) *CBA1* and (D) *SHMT* genes in four (Pt2, Pt3, Pt4 and Pt8) of the ten accessions with the presence of vitamin B12 in axenic cultures (light gray), and with natural bacteria and no vitamin B12 in the growing media (black).

## Discussion

Using whole genome sequence analysis of accessions sampled across multiple geographic locations around the world (Fig. S1), the aim of this study was to describe the global genetic and functional diversity of the model diatom *Phaeodactylum tricornutum*. By defining a comprehensive landscape of natural variations across multiple accessions, we could investigate genetic structure between *P. tricornutum* populations, and a summary of our results is presented in Fig. 7. To do so, we first performed reference-based assembly and found consistently high genome coverage (>90%) mapped by sequencing reads from respective accessions, where some accessions have more coverage (>98%, Pt1, Pt2, Pt3, and Pt9) than others (Table 1). This difference is independent of the size of the sequencing library, as it does not correlate with the genome coverage (Table 1), and a portion of unmapped reads is likely a consequence of the incomplete reference genome, which contains several gaps [1]. Additionally, given the redundant nature of unmapped reads together with the fact that the unmapped reference genome is annotated as being rich in TEs (Fig. S2), a major portion of unmapped reads likely account for large structural variability within the genomes of individual accessions. This explanation is most clear in Pt10, which is shown to have the largest number of gene losses (Fig. 3B) and the highest number of accession-specific TEs with high copy numbers (Fig. 3C) and covers at least (92%) of the reference genome (Table 1). This suggests the role of TEs in creating substantial genetic diversity as also shown in many species of plants and animals [71, 72].

**Figure 7.** Population structure of the ecotypes. The color gradient from yellow to blue indicates low to high numerical values across each ecotype (indicated on top X-axis of panel A) within different functional categories indicated on Y-axis. These functional categories include (from top to bottom), Year of sampling = Year in which the respective ecotype was sampled, Total SNP = Absolute number of SNPs found in each ecotype, Specific/Total SNP (%) = percentage of ecotype specific SNPs, Heterozygosity = Number of heterozygous SNPs from a set of total SNPs within each ecotype, Homozygosity = Number of homozygous SNPs from a set of total SNPs within respective ecotype, Sites deviated from HWE = Number of SNP

Next, based on patterns of variations discovered over the whole genomes and on the molecular marker genes (18S and ITS2) of all the accessions, and by using various clustering algorithms (see Results), the ten accessions could be grouped into four genetic clades. Clade A clusters Pt1, Pt2, Pt3, Pt9; clade B includes Pt4; clade C clusters Pt5, Pt10; and clade D clusters Pt6, Pt7, Pt8. Most of the structural variants discovered, both small (SNPs and INDELS) and large (CNV and Gene Loss), are shared among populations within a clade rather than between clades. This suggests high intra-clade relatedness over a variety of structural, functional and possibly ecological traits.

*P. tricornutum* is a coastal species with limited dispersal potential, which is consistent with the reports of its absence in the open ocean from *Tara* Oceans data [7]. Consequently, the Fixation index (*Fst*) between different genetic clades is very high (0.2 – 0.4) (Fig. 1D), confirming the existence of accessions subdivisions into four genetic clades. As also expected for an organism with limited dispersal potential, the accessions show partial geographical structuring (Fig. 2A, 2B), as Pt5, Pt9 and Pt10 clusters with accessions not sampled from proximal locations. These dispersals to different localities may be fostered by ocean currents [42], human activities like rafting, ballasting [73, 74], and migration of birds [75–77]. In addition, the fact that the subdivisions do not correlate with the sampling time (Fig. 7, Fig. S1), which spans approximately a century, suggests long and stable genetic populations, which is in line with reports from other diatom species [41, 42]. This suggests as reported in these studies that environmental conditions have a more important impact in structuring the populations than dispersal potential and generation time.

Although there exist partial genetic structuring within the accessions, the average nucleotide diversity (π), estimated across all the accessions, is remarkably low (0.2%) compared to the diversity estimates in other unicellular eukaryotes [36, 48, 78–80] but in line with previous estimations in marine phytoplanktonic eukaryotes [81]. Given the observation that there exist a large proportion of heterozygous variant alleles (Fig. 1C), the high *Fst* between the clades, and the low nucleotide diversity across the accessions, we propose that allele frequency plays a significant role in the genetic differentiation of the clades. The difference in allele frequencies is possibly linked to adaptive selection. This phenomenon has recently been studied in diatoms where allele-specific expression of numerous loci has been demonstrated to be a significant source of adaptive evolution in the cold-adapted diatom species *Fragilariopsis cylindrus* [82]. Furthermore, high proportions of heterozygous variant alleles in some clades (clade A, 98%, Fig. 7) compared to others (clade B, 45%, Fig. 7) suggests a high selection pressure in the clade B accession Pt4). High *Fst*, and yet low nucleotide diversity across all the accessions, suggests some degree of genetic and functional convergence among the accessions. This can be explained as a consequence of laboratory culture mediated domestication where some genes accumulate nonsynonymous mutations which might result from the relaxation of selective constraints in an artificial environment.

It is also worth considering that genetic homogenization or functional convergence across the meta-population can also be a consequence of continuous gene flow between the accessions. However, in the case of *P. tricornutum*, gene flow seems limited as highly differentiated populations show partial geographical structure, except Pt9 of clade A, Pt5 of clade C and Pt8 of clade D (Fig. 2A, 2B). This is consistent with earlier findings in *Emilinia huxleyi* and *Thalassiosira pseudonana* in which genomes of isolates from different geographical locations clustered together in the phylogeny [83, 84]. In addition, *P. tricornutum* is not known to reproduce sexually, although various components (genes) of the meiosis pathway are conserved in *P. tricornutum* as well as in other diatom species known to undergo sexual reproduction [85]. Furthermore, the absence of contemporary base changes (CBC) within ITS2 secondary structure between all the accessions compared to the presence of many CBCs between *P. tricornutum* accessions and other diatom species suggests that the accessions may be able to exchange genetic material sexually. However, because *P. tricornutum* is a coastal diatom with only limited dispersal capacity, which is further supported by its apparent absence in the open ocean [7], the possibility of gene flow within different populations is likely to be limited at best.

Next, high linkage disequilibrium (>0.7) observed across all the accessions (Fig. S3B) can be explained by prolonged asexual reproduction [86], a common behavior among diatoms [84]. Asexual reproduction results in higher proportions of divergent alleles within loci with less genetic variation among individuals, and a significant deviation from HWE [86]. Therefore, it is likely that clade B with one isolate, Pt4 also undergoes sexual reproduction reasonably often compared to clade A accessions, as Pt4 possesses the smallest number of heterozygous variant alleles, most of which follow HWE (Fig. 1C, 7). To our surprise, despite high variability in the levels of heterozygosity between different accessions (Fig. 7), the mutational spectrum, compared to the reference, and across all the accessions consisted of high G:C -> A:T and A:T -> G:C transitions (Fig. S3A). Deamination of cytosines dominantly dictates C to T transitions in both plants and animals [87, 88], and CpG methylation potential of the genome is greatly influenced by heterozygous SNPs in CpG dinucleotides [89]. Previous studies have demonstrated low DNA methylation in *P. tricornutum*, using Pt1 8.6, a monoclonal isolate accessioned from a Pt1 single cell as a reference [32, 90]. Because there exist significant differences in the proportion of heterozygote variant alleles between the accessions (45% −98%), testing for DNA methylation patterns across different accessions may provide an interesting opportunity to dissect cross-talk between loss of heterozygosity and DNA methylation in the selection of certain traits [91].

The four genetic clades are further supported by functional specialization of grouped populations (Fig. 5), nicely illustrated with Pt4 in clade B. Pt4 shows a low non-photochemical quenching capacity (NPQ) [18], which was proposed to be an adaptive trait to low light conditions. Specifically, this accession has been proposed to establish an up-regulation of a peculiar light harvesting protein LHCX4 in extended dark conditions [18, 20]. In line with these observations, a gene involved in nitrate assimilation (Phatr3_EG02286) in Pt4 shows high copy numbers, suggesting an altered mode of nutrient acquisition. Nitrate assimilation was shown to be regulated extensively under low light or dark conditions to overcome nitrate limitation of growth in *Thalassiosira weissflogii* [92]. Pt4 is likely adapted to the low light and highly seasonal environment that characterizes the high latitudes where it was found, which may well affect its nitrate assimilation capacity [93, 94]. Additional functions emerging from clade C (Pt5 and Pt10) include vacuolar sorting and vesicle-mediated transport gene-families, which could be an indication of altered intracellular trafficking [95].

In conclusion, the study presents pan-genomic diversity of the model diatom *P. tricornutum*. This is the first study within diatoms that provides a comprehensive landscape of diversity at whole genome sequence level and brings new insights to our understanding of diatom functional ecology and evolution. Given our observation that *P. triconrutum* accessions possess high numbers of heterozygous alleles, it would be interesting to think of possible selective functional preferences of one allele over the other under different environmental conditions or during the life/cell cycle. In the future, such studies could be crucial for deciphering the mechanisms underpinning allele divergence and selection within diatoms. Likewise, more than answers, our study delivers more questions, which should help address the genetic basis of diatom success in diverse ocean ecosystems. Finally, this study provides the community with genomic sequences of *P. tricornutum* accessions that can be useful for functional studies.

## Experimental procedures

### Sample preparation, sequencing and mapping

Ten different accessions of *P. tricornutum* were obtained from the culture collections of the Provasoli-Guillard National Center for Culture of Marine Phytoplankton (CCMP, Pt1=CCMP632, Pt5=CCMP630, Pt6=CCMP631, Pt7=CCMP1327, Pt9=CCMP633), the Culture Collection of Algae and Protozoa (CCAP, Pt2=CCAP 1052/1A, Pt3= CCAP 1052/1B, Pt4= CCAP 1052/6), the Canadian Center for the Culture of Microorganisms (CCCM, Pt8=NEPCC 640), and the Microalgae Culture Collection of Qingdao University (MACC, Pt10=MACC B228). All of the accessions were grown axenically in batch cultures with a photon fluency rate of 75 μmol photons m-2 s-1 provided by cool-white fluorescent tubes in a 12:12 light: dark (L:D) photoperiod at 20 °C. Exponentially growing cells were harvested and total DNA was extracted with the cetyltrimethylammonium bromide (CTAB) method[96]. At least 6 μg of genomic DNA from each accession was used to construct a sequencing library following the manufacturer’s instructions (Illumina Inc.). Paired-end sequencing libraries with a read size of 100 bp and an insert size of approximately 400 bp were sequenced on an Illumina HiSeq 2000 sequencer at Berry Genomics Company (China). The corresponding data can be accessed using bioSample accessions: SAMN08369620, SAMN08369621, SAMN08369622, SAMN08369623, SAMN08369624, SAMN08369625, SAMN08369626, SAMN08369627, SAMN08369628, SAMN08369629, SAMN12551644 (Pt3Ov genomic), SAMN12551645 (Pt3Ov Transcriptomic), and SAMN12551646 (Pt8Tc genomic). Low quality read-pairs were discarded using FASTQC with a read quality (Phred score) cutoff of 30. Using the genome assembly published in 2008 as reference [1], we performed reference-assisted assembly of all the accessions. We used BOWTIE (-n 2 –X 400) for mapping the high quality NGS reads to the reference genome followed by the processing and filtering of the alignments using SAMTOOLS and BEDTOOLS. Detailed methods are provided in File S4.

### Discovery of small polymorphisms and large structural variants

GATK [46], configured for diploid genomes, was used for variant calling, which included single nucleotide polymorphisms (SNPs), small insertions (of varying lengths from 1 bp to 312 bp) and deletions (of lengths from 1 bp to 400 bp). The genotyping mode was kept default (genotyping mode = DISCOVERY), Emission confidence threshold (-stand_emit_conf) was kept 10 and calling confidence threshold (-stand_call_conf) was kept at 30. The minimum number of reads per base, to be called as a high quality SNV, was kept to 4 (read-depth >=4x). Following this filtration step, the number of sites in the protein coding genes covered for all 10 accessions, and therefore callable to estimate the genome wide synonymous and non-synonymous polymorphism, added up 11.0 Mbp. The average pairwise synonymous and non-synonymous diversity π_S_ and π_N_ [97] were estimated for all genes using in-house R script from [97] equation 22 for each gene and the complete callable coding sequences (available from the authors upon request).

Next, considering Z-score as a normalized measure of read-depth, gene and TE candidates showing multiple copies (representing CNV) or apparently being lost (representing gene loss) were determined. For TE CNV analysis, TEs that are more than 100 bp lengths were considered. We measured the fold-change (*Fc*) by dividing normalized read depth per genomic feature (Z-score per gene or TE) by average of normalized read depth of all the genes/TEs (average Z-score), per sample. Genes or TEs with log2 scaled fold change >=2 were reported and considered to exist in more than one copy in the genome. Genes where the reads from individual accession sequencing library failed to map on the reference genome were considered as potentially lost within that accession and reported. Detailed method is provided in File S4. Later, some randomly chosen loci were picked and validated for the loss in the accessions compared to the reference genome by PCR analysis.

### Validation of gene loss and quantitative PCR analysis

In order to validate gene loss, DNA was extracted from all the accessions as described previously [22] and PCR was performed with the primers listed in Table S1. PCR products were loaded in 1% agarose gel and after migration gels were exposed to UV light and photographs were taken using a gel documentation apparatus to visualize the presence and absence of amplified fragment. To assess gene expression, RNA was extracted as described in [23] from three biological replicates of accessions grown axenically in Artificial Sea Water (ASW) [98] supplemented with vitamins as well as in the presence of their endemic bacteria in ASW without vitamins. qPCR was performed as described previously [23]. Briefly, cDNA was synthetized from 1 µg RNA using High-Capacity cDNA Reverse Transcription Kit (catalogue number 4368813) from Fischer scientific and according to manufacturer instructions. 1µl of cDNA was used in the QPCR reaction with the LightCycler^®^ 480 SYBR Green I Master (catalogue number 04707516001) from Roche and according to manufacturer instructions. Two reference genes were used, Tata Box binding Protein and Ribosomal Protein Small subunit 30S for normalization [23].

### *P. tricornutum* population structure

#### Haplotype analysis

First, to cluster the accessions as haplogroups, ITS2 gene (chr13: 42150-43145) and 18S gene (chr13: 43553-45338) were used. Polymorphic sites across all the accessions within ITS2 and 18S genes were called and used to generate their corresponding accession specific sequences, which were then aligned using CLUSTALW. The same approach was employed to perform haplotype analysis at the whole genome scale. Later, a maximum likelihood algorithm was used to generate the 18S, ITS2 and, whole genome tree with bootstrap values of 1,000. We used MEGA7 [99] to align and deduce the phylogenetic trees.

#### CBC analysis

CBC analysis was done by generating the secondary structure of ITS2 sequences, using RNAfold [100], across all *P. tricornutum* accessions and other diatom species. The other species include one centric diatom species *Cyclotella meneghiniana* (AY906805.1), and three pennate diatoms *Pseudo-nitzschia delicatissima* (EU478789.1), *Pseudo-nitzschia multiseries* (DQ062664.1), *Fragilariopsis cylindrus* (EF660056.1). The centroid secondary structures of ITS2 gene with lowest minimum free energy were used for CBC analysis. We used 4SALE [101] for estimating the presence of CBCs between the secondary structure of ITS2 gene across all the species.

#### Population genetics

Further, we measured various population genetic functions to estimate the effect of evolutionary pressure in shaping the diversity and resemblance between different accession populations. Within individual accessions, by using approximate allelic depths of reference/alternate alleles, we calculated the alleles that are deviated from Hardy Weinberg equilibrium (HWE). We used chi-square estimation to evaluate alleles observed to deviate significantly (P-value < 0.05) from the expected proportion as per [p2 (homozygous) + 2pq (heterozygous) +q2 (homozygous) =1) and should be 0.25% + 0.50% + 0.25%. Alleles were considered heterozygous if the proportion of ref/alt allele is between 20-80%. The proportion of ref/alt allele was calculated by dividing the number of reads supporting ref/alt base change by total number of reads mapped at the position. We evaluated average R^2^ as a function to measure the linkage disequilibrium with increasing distance (1 kb, 5 kb, 10 kb, 20 kb, 30 kb, 40 kb and 50 kb) between any given pair of mutant alleles across all the accessions using expectation-maximization (EM) algorithm deployed in the VCFtools. Although no recombination was observed within the accessions, attempts were made to look for recombination signals using LDhat [102] and RAT [103]. Genetic differentiation or variability between the accessions was further assessed using the mathematical function of Fixation index (FST), as described by Weir and Cockerham 1984 [104].

#### Genetic clustering

Genetic clustering of the accessions was done using Bayesian clustering approach by applying *Markov Chain Monte Carlo* (MCMC) estimation programmed within ADMIXTURE (version linux-1.3.0) [50]. Accessory tools like PLINK (version 1.07-x86_64) [105] and VCFtools (version 0.1.13) [106] were used to format the VCF files to ADMIXTURE accepted formats. In the absence of data from individuals of each accession/sample, we assumed the behavior of each individual in a sample to be coherent. Conclusively, instead of estimating the genetic structure within an accession, we compared it across all the accessions. We first estimated the possible clusters of genomes, (K), across all the accessions, by using cross-validation error (CV error) function of ADMIXTURE [107]. We chose the value of K with lowest cross-validation error (see extended methods, File S4). Finally, we used ADMIXTURE with 200 bootstraps, to estimate the genome clusters within individual accessions by considering the possible number of genomes derived via CV-error function.

### Functional characterization of polymorphisms

snpEff [108] and KaKs [109] calculator were used to annotate the functional nature of the polymorphisms. Along with the non-synonymous, synonymous, loss-of-function (LoF) alleles, transition to transversion ratio and mutational spectrum of the single nucleotide polymorphisms were also measured. π_N_/π_S_ ratios were calculated for 5232 protein coding genes containing more than 10 SNP. 10% of genes with lower π_N_/π_S_ were considered as under strong purifying selection on amino-acid composition (File S3). Genes with π_N_/π_S_ >1, and average frequency on non-synonymous polymorphism higher than the average frequency of synonymous polymorphism were considered as candidate genes under balancing selection on amino-acid composition (File S3). Various in-house scripts were also used at different levels for analysis and for plotting graphs. Data visualization and graphical analysis were performed principally using ClicO [110], CYTOSCAPE [111], IGV [112] and R (https://www.r-project.org/about.html). Based on the presence of functional domains all the Phatr3 genes [63] were grouped into 3,020 gene families. Subsequently, the constituents of each gene family were checked for being either affected by loss-of-function mutations or under relaxed selective constraints. To estimate an unbiased effect of any evolutionary pressure (LoF allele or balancing selection mutations) on different gene families, induced because of high functional redundancies in the gene families, a normalized ratio named as effect ratio (EfR), was calculated. Precisely, the EfR normalizes the fact that if any gene family have enough candidates to buffer the effect on some genes influencing evolutionary pressures, it will be considered as less affected compared to the situation where all or most of the constituents are under selection pressure. The ratio was estimated as shown below and gene families with EfR larger than 1 were considered as being significantly affected.

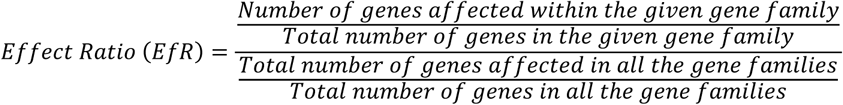

Additionally, significantly enriched (chi-square test, P-value < 0.05) biological processes associated within genes experiencing LoF mutations, purifying selection, balancing selection (BS), or showing CNV, or being lost (GnL), were estimated by calculating observed to expected ratio of their percent occurrence within the given functional set (BS, LoF, CNV) and their occurrence in the complete annotated Phatr3 (http://protists.ensembl.org/Phaeodactylum_tricornutum/Info/Index) biological process catalog. Later, considering gene family EfR as a function to measure the association rate, we deduced Pearson pairwise correlations between different accessions. The correlation matrix describes that if many equally affected gene families are shared between any given pair of accessions, they will have higher correlation compared to others. Finally, hierarchical clustering using Pearson pairwise correlation matrix assessed the association between the accessions.

## Acknowledgements

HH acknowledges support from National Natural Science Foundation of China (grant No. 91751117). GW acknowledges the Strategic Priority Research Program of the Chinese Academy of Sciences (grant No. XDA17010502). CB acknowledges funding from the ERC Advanced Award ‘Diatomite’, the LouisD Foundation of the Institut de France, the Gordon and Betty Moore Foundation, and the French Government ‘Investissements d’Avenir’ programmes MEMO LIFE (ANR-10-LABX-54), PSL* Research University (ANR-1253 11-IDEX-0001-02), and OCEANOMICS (ANR-11-BTBR-0008). CB also thanks the Radcliffe Institute of Advanced Study at Harvard University for a scholar’s fellowship during the 2016-2017 academic year. LT acknowledges funds from the CNRS, MEMO LIFE (ANR-10-LABX-54) and the region of Pays de la Loire (ConnecTalent EPIALG project). AR was supported by an International PhD fellowship from MEMO LIFE (ANR-10-LABX-54).

## Contributions

LT, HH and CB conceived the study. LT, AR and GP designed the study. GW, PV, AFDC, CC and LT did the experiments. AR, FRJV, AV and GP developed and performed the bioinformatics analysis. AR, GP, FRJV and LT interpreted the results. AR and LT wrote the manuscript with input from all the authors. LT coordinated and supervised the study.

